# Seq2scFv: a toolkit for the comprehensive analysis of display libraries from long-read sequencing platforms

**DOI:** 10.1101/2024.07.04.602016

**Authors:** Marianne Bachmann Salvy, Luca Santuari, Emanuel Schmid-Siegert, Nikolaos Lykoskoufis, Ioannis Xenarios, Bulak Arpat

## Abstract

Antibodies have emerged as the leading class of biotherapeutics, yet traditional screening methods face significant time and resource challenges in identifying lead candidates. Integrating highthroughput sequencing with computational approaches marks a pivotal advancement in antibody discovery, expanding the antibody space to explore. In this context, a major breakthrough has been the full-length sequencing of single-chain variable fragments (scFvs) used in *in vitro* display libraries. However, few tools address the task of annotating the paired heavy and light chain variable domains (VH and VL), which is the primary advantage of full-scFv sequencing. To address this methodological gap, we introduce Seq2scFv, a novel open-source toolkit designed for analyzing *in vitro* display libraries from long-read sequencing platforms. Seq2scFv facilitates the identification and thorough characterization of V(D)J recombination in both VH and VL regions. In addition to providing annotated scFvs, translated sequences and numbered chains, Seq2scFv enables linker inference and characterization, sequence encoding with unique identifiers and quantification of identical sequences across selection rounds, thereby simplifying enrichment identification. With its versatile and standalone functionality, we anticipate that the implementation of Seq2scFv tools in antibody discovery pipelines will efficiently expedite the full characterization of display libraries and potentially facilitate the identification of high-affinity antibody candidates.

## 1 Introduction

Due to their high target specificity and strong binding affinity, antibodies currently stand as the leading class of biotherapeutics and their market share is continuously expanding [27]. However, traditional antibody screening methods, although efficient and well-established, are cumbersome, time-consuming and technically challenging. Moreover, they are limited to the analysis of a small fraction of the antibodies generated. Using experimental approaches alone, it remains difficult to identify binders aimed at different epitopes of the target and with a variety of biophysical characteristics, including properties relevant for their developability [30] [13] [19].

Incorporating computational approaches and Next Generation Sequencing (NGS) into antibody development pipelines for rational design can effectively overcome these limitations. Indeed, NGS-based computational methods can track the progressive enrichment of target-specific sequences in *in vitro* display experiments, guiding the selection of lead antibody candidates with higher sequence and epitope diversity [35] [36] [4]. Furthermore, computational techniques and novel Artificial Intelligence (AI) methods prove valuable in later stages of antibody development, including improvement of antibody binding affinity, humanization and homology modelling by leveraging structural bioinformatics or biophysical property characterization [8] [7] [31] [19].

Unlocking the full potential of computational and AI methods in immunogenetics relies on accurately identifying and thoroughly characterizing antibody sequences [39]. This process typically begins with mapping the reads against germline gene databases to identify V(D)J rearrangements in each Vdomain, with tools such as IgBLAST [43], IMGT/HighV QUEST [26], MiXCR [5] and many more, extensively reviewed in Norman *et al*., 2020 [31] as well as Smakaj *et al*., 2020 [39]. These tools generally also identify the correct translation frame of each variable domain (V-domains) and delimit their V, D and J regions. Further characterization involves antibody numbering, which assigns numerical identifiers to individual amino acid residues within the V-domains. This contextualizes each position within the antibody structure according to established schemes such as Chothia [6], Kabat [18], IMGT [23] and several others reviewed in Dondelinger *et al*., 2018 [9]. Tools such as ANARCI [10], AbRSA [25], AbNum [1] or AntPack [34] have been developed for antibody numbering.

In parallel to advancements in bioinformatics methods, the innovative application of Pacific Biosciences’ (PacBio) long-read HiFi sequencing to *in vitro* display libraries has enabled the high-throughput full-length sequencing of recombinant antibodies. Previously, reading both the heavy (VH) and light

(VL) variable domains that constitute single chain variable fragments (scFvs) was a challenge due to the limitation of most NGS platforms to fragments of 500 bp, while scFvs typically span around 850 base pairs (bp) [15]. Fully sequencing both chains is particularly valuable. Indeed, while the complementarity determining regions (CDRs) are known to play a significant role in binding [2], with the third CDR of the heavy chain (HCDR3) being crucial for antibody specificity [42], evidence suggests that other regions also contribute to binding [20]. Furthermore, the light chain plays a role in specificity [11], with specific regions influencing the structure of the HCDR3 loop [17]. Therefore, sequencing the entire scFv provides crucial information for identifying better binders [24]. The advent of Single Molecule Real-Time (SMRT) sequencing on PacBio platforms has already enabled the identification of antibodies with a diverse range of affinities, epitopes and biophysical characteristics, by leveraging full-length scFvs information in *in vitro* display libraries [30] [12].

Currently, there is a gap in the landscape of antibody annotation methods as not many tools address the annotation of VH and VL in a single read, a requisite for the analysis of full length scFvs. Though IMGT/HighV-Quest [26] does provide a scFv analysis option, it is only provided as a web server and is not open source. Additionally, it limits the V(D)J alignment to IMGT sequences, as it does not provide the option of using custom germline gene databases.

Here, we introduce Seq2scFv, a comprehensive collection of integrated approaches and scripts tailored for analyzing *in vitro* display libraries derived from long-read sequencing. Open-source for academic use under a non-commercial license, while subject to a separate license for any other purposes, this toolkit streamlines widely used bioinformatic tools with a suite of custom algorithms to process sequences into fully delimited and annotated scFvs. Scripts are provided for sequence cataloguing and encoding, characterization and numbering of both V-domains, scFv delimitation and conformation validation. Additionally, a framework for the inference of the linker sequence from the data is provided, together with scripts for evaluating the most frequent linker sequences and their length distribution. The comparison between the consensus or reference linker sequence (if known beforehand) and the linker sequence in each individual read can be used as an additional quality parameter for selecting well-annotated scFvs. A script for flagging scFvs on other quality criteria is also provided. Importantly, Seq2scFv can quantify representative sequences across successive selection rounds, facilitating downstream enrichment analysis for frequency-based lead antibody candidate selection. Outputs from the different analysis steps in the framework are primed for easy analysis, sub-selection, or exportation to human-readable formats. Emphasizing flexibility and consistency, this collection of tools accommodates user-specified parameters, including IMGT and Kabat annotation and numbering schemes, as well as user-provided germline gene databases.

In addition to introducing Seq2scFv, we demonstrate its real-world application using openly accessible data from Nannini and colleagues, published in 2020 [30]. By processing this example dataset, we showcase the use of Seq2scFv in analyzing *in vitro* display libraries derived from long-read sequencing platforms. Our aim is to provide researchers with a practical toolkit for comprehensive antibody characterization. It is outside the scope of this article to select potential binding candidates based on Seq2scFv annotation and experimentally validate them. For validation of the NGS-guided selection as a frequency-based approach to identify potential binders, we refer to the original publication by Nannini *et al*. [30] and the growing literature in the field [12] [35] [36] [4] [29]. Beyond its application in antibody discovery, characterized scFvs can be compiled in large datasets to leverage machine learning applications for antibody discovery and engineering [24].

## 2 Methods

The main steps for analyzing long-read display libraries with Seq2scFv consist in tagging sequences with unique IDs, library merging, V(D)J alignment, scFv delimitation, linker detection and scoring, flagging and counting. Figure 1 provides an overview of the analysis framework, and the sections below describe in detail the tools developed and implemented, as well as the type and origin of the publicly available dataset.

**Figure 1.**
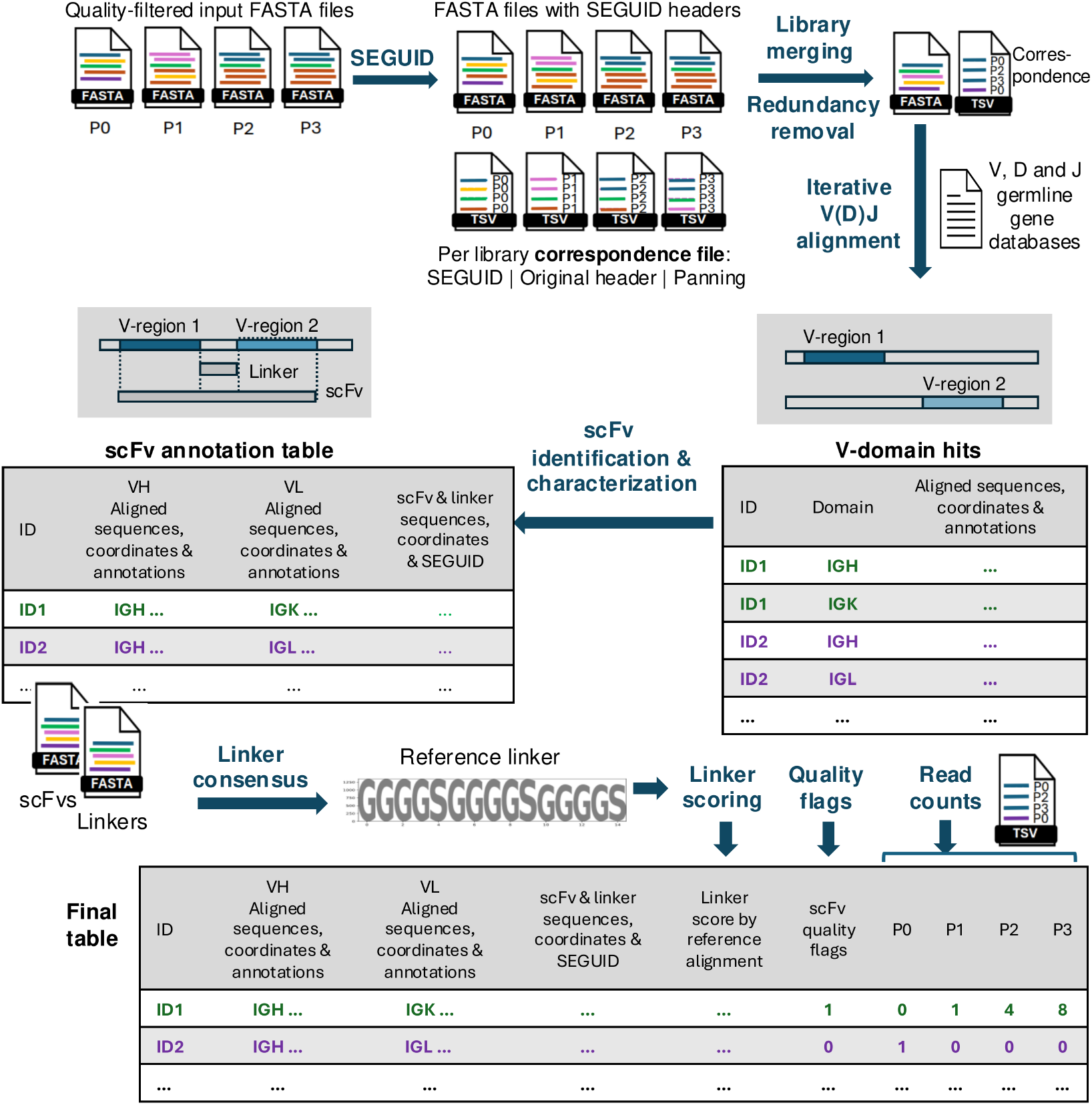
The main processes are indicated with blue arrows and text. Input FASTA files from different panning rounds (P0, P1, P2, P3) undergo renaming with SEGUIDs. They are then merged, cataloged, and deduplicated to remove sequence redundancy. Unique reads are aligned to V, D, and J regions, resulting in a list of VH and VL hits. Reads with one VH and one VL are selected for further scFv delineation, linker sequence inference, and evaluation. The process of identifying scFvs from the alignment hits is illustrated in the grey boxes. The final output includes quality flags and read count data across panning rounds.

### 2.1 Input, library merging and SEGUID cataloguing

FASTA files, each representing a distinct library sample (or “panning round”), are provided as input. It is expected that demultiplexing, adapter trimming, and reverse-complementation (to ensure all sequences are in the same orientation) have already been performed. Additional quality preprocessing can also include filtering out sequences falling outside of the expected insert size interval. In subsection 2.7.2, we detail the preprocessing methods used to prepare the example dataset.

To facilitate the tracking of clone amplification through panning rounds, all sequences from the different panning rounds are pooled into a single FASTA file before antibody identification and characterization. This process involves assigning a Sequence Globally Unique Identifier (SEGUID; [3]) to each sequence and maintaining a correspondence file to track its library origin. Using seqkit v2.7.0 [37], only one representative among identical sequences encoded by the same SEGUID is retained in the combined FASTA file, effectively removing redundancy and optimizing the computational workload for downstream antibody characterization. Despite this, the correspondence file preserves the occurrence of sequences across different panning rounds, ensuring that individual clone amplification monitoring is not compromised.

### 2.2 ALIGNMENT of sequences to germline V, D and J genes

The identification of VH and VL in the query read employs a method resembling the IMGT/HighVQUEST algorithm for scFv sequence analysis [14]. In IMGT/HighV-QUEST, a V-domain is initially identified and characterized, followed by the search and characterization of a second V-domain in the flanking regions, provided they exceed a length threshold of 150 bp. Then, coordinates of the two V-domains are used to delimit the position of the linker sequence.

In this approach, we employ IgBLAST [43], a specialized alignment tool for immunoglobulins, to identify V-domains. By mapping the nucleotide sequence against a user-provided germline gene database, IgBLAST delineates the V, (D) and J regions that constitute a V-domain, generating a tabular file conforming to the Adaptive Immune Receptor Repertoire (AIRR) Rearrangement format [41]. The table encompasses matched germline genes, nucleotide and amino acid alignment sequences, coordinates and scores, and framework region (FWR) and CDR sequences and coordinates (further details provided in the Supplementary table 2). IgBLAST v. 1.20.0 is used in an iterative process in a Python script: once a V-domain is identified, adjacent regions are split, and if they meet a user-defined length threshold, they are submitted to a new IgBLAST search. This iterative process annotates all possible V-regions within a read until no additional splitting is possible. Finally, the script reunites all V-domain hits for all sequences into a single tab-separated file, with one hit per row, and adjusts the alignment coordinates to reflect the original sequence.

### 2.3 scFv delimitation and characterization

To identify single-chain variable fragments (scFvs), a custom set of Python scripts are used. Initially, reads with precisely one VH (IGH locus) and one VL (IGK or IGL locus) hit reported by IgBLAST are selected, while sequences with a different number of hits are excluded. Columns are annotated with either VH or VL prefixes according to the identified locus. Subsequently, the VH and VL information is merged, consolidating each row in the table to represent a putative scFv rather than individual hits (similar to the format for paired antibodies used in the Observed Antibody Space database [32]).Additionally, sequences containing a stop codon in either VH or VL domain, as determined by IgBLAST’s sequence translation, are removed from further consideration.

Next, the translation frame of the full read is determined, ensuring that both VH and VL amino acid sequence alignments are contained within the same frame. Sequences failing to meet this criterion are designated as “out of frame”. Upon identifying the read’s translation frame, the scFv is delineated at both the amino acid and nucleotide levels leveraging IgBLAST’s alignment coordinates. The start and end of the scFv are defined by the 5’ end of the first V-domain and the 3’ end of the second V-domain, respectively, while the linker region is bounded by the 3’ end of the first V-domain and the 5’ end of the second V-domain.

Based on these delimitations, sequences and positions of the scFv and linker at both the nucleotide and amino acid levels are incorporated into the table. Additionally, scFv sequences are encoded with a SEGUID to facilitate clone cross-library amplification and cross-experiment comparison. Furthermore, scFv characterization by Seq2scFv tools involves numbering of both V-domains using AntPack, which employs a global alignment approach with a custom scoring matrix [34]. Alongside a list of numbered positions and other AntPack annotations, fields reporting the consistency between alignment and numbering approaches are included (further details provided in Supplementary table 2).

The final steps of scFv characterization with Seq2scFv tools involve annotating CDR and FWR regions. While IgBLAST reports sequences at both nucleotide and amino acid levels, positions are only provided at the nucleotide level and are relative to the full read. To simplify integration with downstream analyses, amino acid and nucleotide coordinates are added and updated, respectively, to be relative to the corresponding delimited chain.

After completing the annotation process, output tables and FASTA files are generated. The FASTA files contain nucleotide and amino acid sequences of scFvs and linkers, while the tables report the annotations. An ‘in-frame’, paired, and delimited table consolidates all correctly annotated scFvs, while three separate tables compile unpaired V-domains, scFvs with non-standard amino acids, and out-of-frame scFvs.

### 2.4 Linker sequence evaluation

The characterization of scFvs by Seq2scFv can include a systematic and thorough assessment of the inferred linker sequences, which may deviate from the designed linker due to mutations or annotation issues. For instance, if an uncommon V or J gene is present in one of the V-domains, or somatic hypermutation (SMH) introduced many mutations, the alignment might fail to accurately pinpoint its position in the read and infer the correct start and end coordinates of the linker sequence. Reporting linker sequence discrepancies enables a more comprehensive assessment of annotation quality.

Identifying linker sequence divergences requires an amino acid reference linker, which can either be supplied by the user or inferred from the data. To derive a consensus sequence from the dataset, nucleotide and amino acid linkers files are randomly subsampled from their respective FASTA files and aligned with Clustal Omega [38]. Subsequently, sequence logos are generated using LogoMaker [40], along with tables reporting the weight (fraction of non-gapped symbols) of each position in the alignments. This weight data enables the determination of the most prevalent nucleotides or amino acids in significant positions, defined by exceeding a predetermined weight threshold. In instances where no user-provided reference linker is available, the consensus linker is considered as reference. Additionally, to provide further characterization of the linkers, tools in Seq2scFv can be used to generate sequence logos and a report containing the top ten linkers ranked by frequency, along with frequency tables and histograms depicting linker lengths.

Furthermore, the scFv annotation table is revisited to align each reported linker with the reference linker. This alignment process, incorporating customized penalties to enhance alignment score interpretability (for more details, refer to Seq2scFv documentation), is conducted using the SmithWaterman algorithm. Three columns are added to provide insights into linker annotation: the percentage of aligned amino acids relative to the length of the reference linker sequence, the number of mismatched positions, and the count of non-aligned amino acids in the inferred linkers, referred to as “overhang.”

### 2.5 scFv quality flags

Quality flags can be added with tools in Seq2scFv tools to facilitate downstream analysis and filtering based on predefined criteria. These flags utilize a binary annotation (0/1) to indicate whether a given scFv meets specific quality thresholds. Criteria assessed include the correct order of VH and VL domains as well as the linker annotation quality, defined by the user by providing thresholds for the alignment score, the number of mismatches and the length of the linker overhang. Additionally, flags are assigned based on whether each variable domain exceeds a minimum amino acid sequence length and meets IgBLAST criteria for being classified as “productive” (i.e., the V(D)J rearrangement frame is in frame, no stop codon is present, and there are no internal frame shifts in the V gene) and “complete” (i.e., the sequence alignment spans the entire V(D)J region from the first V gene codon to the last complete codon of the J gene).

### 2.6 Read counts

The final process in the framework proposed by Seq2scFv generates count data for each read listed in the scFv annotation table. This process retrieves entries from the correspondence file, where all identical reads were encoded with a SEGUID, while also recording their originating libraries. Thus, for each SEGUID-encoded read in the annotation table, one column per panning round is appended. These columns denote the frequency with which identical read was observed in each library. Leveraging the comprehensive annotations in the scFv annotation table, these counts can be aggregated according to various regions of interest, such as full amino acid or nucleotide scFv sequences, VH or VL domains, CDRs, among others.

### 2.7 Example dataset

#### 2.7.1 Phage libraries

The phage libraries utilized to showcase the application of Seq2scFv were sourced from the publication by Nannini and colleagues (2020) [30]. In their study, phage display libraries were generated through immunization of Wistar rats with the CD160 antigen, followed by three iterative biopanning rounds against the targeted gene (further details regarding this study can be found in their publication). The initial phage library (P0), along with the subsequent panning rounds (P1, P2, P3), were sequenced using a Pacific Biosciences RS sequencer. The site https://rdr.ucl.ac.uk/ndownloader/files/28624629 was last accessed in October 2022 to download the published FASTQ files.

#### 2.7.2 Sequence preprocessing

Basseto v. 0.3.7 [33] was used to selectively retain reads with an expected accuracy of 99%. Next, primer sequences used for phage library amplification prior to sequencing were trimmed using cutadapt v4.1 [28], while ensuring orientation consistency through reverse-complementation. Only sequences flanked by the primers detailed in Nannini *et al*. [30]) were kept (5’ primer GTCGTCTTTCCAGACGTTAGT and 3’ primer CAGGAAACAGCTATGAC). Sequences were then further filtered with cutadapt to select those falling within the expected insert size interval of 900-1200 bp. Processed sequences were then transformed into FASTA format using basseto v. 0.3.7. to conform to Seq2scFv input requirements.

#### 2.7.3 Germline gene database

Sequences from Nannini and colleague’s publication (2020) [30] were searched against *Rattus norvegicus* immunoglobulin germline gene sequences. These were obtained from the IMGT reference directory [22]; available at https://www.imgt.org/vquest/refseqh.html) and formatted according to IgBLAST web instructions (available at https://ncbi.github.io/igblast/cook/How-to-set-up.html). The auxiliary file, containing additional information for each J gene to facilitate CDR3 annotation, was obtained from IgBLAST directory, as they provide this file for all IMGT and NCBI databases.

#### 2.7.4 Parameters

For the V(D)J alignment, parameters included the specification of “rat” as the organism, the path to the germline gene databases, a minimum length of 150 bp for the iterative IgBLAST search and a minimum E-score of 0.01 to validate a hit. Subsequently, during the flag addition step, criteria such as a 90% percent identity score and a maximum of 2 mismatches between inferred linkers for each scFv and the consensus linker derived from the data were incorporated. Also, a minimum domain length of 80 base pairs for both VH and VL domains was imposed. Since information regarding the reference linker sequence was absent, the linker parameter was marked as “undefined”. Additionally, for the identification of the consensus linker sequence from the scFvs, a subsampling of 10% of the linker sequences was chosen.

## 3 Results

Full scFv characterization of Nannini *et al*. phage display libraries (2020) [30] was swiftly conducted using the framework and comprehensive collection of tools Seq2scFv. The raw sequence counts from panning rounds 0, 1, 2, and 3 were 15,395; 10,868; 30,607; and 47,013 reads, respectively. After quality filtering and redundancy removal, the 56,953 reads led to the identification of a total of 14,764 unique scFv across all panning rounds after alignment against the *R. norvegicus* immunoglobulin germline genes, scFv delimitation and characterization.

### 3.1 Preprocessing statistics

Records of the number of reads per panning library across preprocessing stages are presented in Table 1. In the case of Nannini *et al*. 2020 [30], publicly available sequences from panning rounds 0, 1, and 2 already exceeded the threshold of a read quality higher than 0.99, while 1,775 sequences from panning round 3 were discarded due to lower quality (3.78%). Between 87.54% and 99.01% of the quality-filtered sequences were correctly flanked by the provided sequences used for PCR amplification. However, due to size filtering, a significant number of reads from panning rounds 0 and 3 were removed, with only 58.16% and 33.67% of the high-quality reads respectively meeting the criteria of correct adapter flanking and an insert size between 900 and 1200 bp. In contrast, the filtering was less stringent for panning rounds 1 and 2 (91.31% and 85.57% respectively). Regarding sequence redundancy, the unselected library (panning round 0) exhibited the highest diversity, with no repeated sequences, while the proportion of identical sequences increased for panning rounds 1 and 2. However, it decreased again for panning round 3.

**Table 1.** Read counts across preprocessing steps.

| Panning | Raw reads | Quality trimmed reads | Quality filtered reads (% of raw reads) | Adapter-trimmed reads | Reads with adapter (% of quality-filtered reads) | Size filtered reads | Reads with adapter and selected size (% of quality-filtered reads) | Unique reads | Unique reads (% of quality, adapter and size filtered reads) |
| --- | --- | --- | --- | --- | --- | --- | --- | --- | --- |
| 0 | 15,395 | 15,395 | 100.00 | 15,243 | 99.01 | 8,953 | 58.16 | 8,953 | 58.16 |
| 1 | 10,868 | 10,868 | 100.00 | 10,470 | 96.34 | 9,924 | 91.31 | 9,889 | 90.99 |
| 2 | 30,607 | 30,607 | 100.00 | 29,137 | 95.20 | 26,283 | 85.87 | 23,574 | 77.02 |
| 3 | 47,013 | 45,238 | 96.22 | 39,600 | 87.54 | 15,232 | 33.67 | 14,893 | 32.92 |

Read counts across preprocessing steps for panning rounds 0 to 3, with percentages of sequences kept relative to the previous processing point.

Additional quality assessment can be achieved by examining the distribution of read lengths. In Figure 2.a., the presence of sequences falling outside the 900 – 1200 bp range may indicate contamination or unpaired VH or VL. Furthermore, narrowing the focus to the specific interval of expected read lenghts (Figure 2.b.) allows the identification of an accumulation of sequences at particular lengths concurrent with an increase in selective pressure (i.e., with successive rounds of panning), implying an amplification of clones with specific lengths and, ultimately, binding affinities.

**Figure 2.**
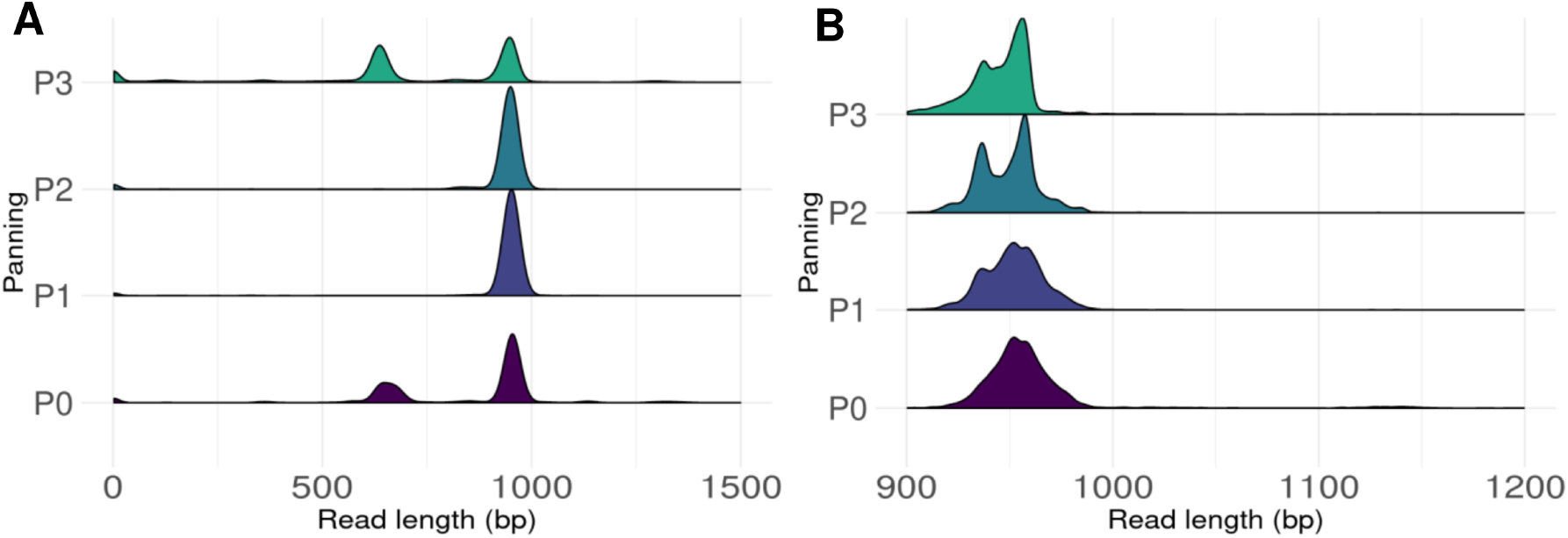
Read length distribution of sequences flanked by the PCR primer sequences. Panel a) displays the distribution without limits, while panel b) exhibits the distribution within the user-provided interval.

### 3.2 scFv characterization

Preprocessing, redundancy removal, SEGUID encoding and pooling of libraries, resulted in a single file containing 56,953 unique sequences. These sequences underwent V-domain searches using IgBLAST, which identified a total of 112,259 V-domains. However, 1,602 V-domains were excluded from further analysis due to hits with E-scores lower than 0.01 at the V or J gene alignments, or because of the absence of a second V-domain within the same read. Additionally, 40,565 sequences were discarded due to being out of frame, either from a discrepancy in frame between the VH and the VL (3,486) or the presence of a stop codon in the linker or any of the V-domains (37,079). However, none of the sequences presented non-standard amino acids within the sequence.

A comprehensive characterization of each domain was compiled into a table format for 14,764 in-frame, paired scFvs. This entailed IgBLAST analysis for each VH and VL, detailing V(D)J gene alignment specifics such as top germline gene match, alignment positions, translations, and FWR and CDR delimitation. Notably, FWR and CDR information was configured to indicate nucleotide and amino acid coordinates relative to each V-domain, using a 1-based system, rather than relative to the full read. Further V-domain annotation included the generation of ungapped versions of VH and VL nucleotide sequences, along with IMGT numbering for each amino acid chain, and assessing alignment versus numbering consistency.

In addition to V-domain annotation, the table also compiled scFv-level and read-level information. This encompassed scFv and linker coordinates relative to the read, as well as their full sequences and SEGUIDs encoding scFv nucleotide and amino acid sequences. Simultaneously, 13,966 unique nucleotide scFv sequences and 13,598 unique amino acid scFv sequences were written to FASTA files, with SEGUIDs serving as headers. Additional columns described the alignment quality of each detected linker against the inferred consensus, referring to the percentage alignment, the mismatched positions, and the alignment overhang. Quality across several user-specified parameters (minimum V-domain lengths, linker annotation quality, scFv conformation) were summarized by different flags, included in the annotation table. Finally, read occurrences across the different panning libraries were reported in the last columns, with one column per panning round.

### 3.3 Linker inference and evaluation

A comprehensive evaluation of inferred linker sequences was undertaken with Seq2scFv. Since no user-supplied linker sequence was available, the consensus amino acid linker sequence (*GGGGS*)_3_ was used as reference for alignment against all other linkers. This consensus linker was derived through alignment and consensus determination from randomly subsampled amino acid linker sequences using Clustal Omega and LogoMaker, respectively. The generated amino acid sequence logo (Figure 3) provides a succint overview of the conservation within the inferred linker sequence, while Table 2 presents the top 10 most frequent amino acid linker sequences identified in the dataset. Furthermore, the distribution of amino acid linker lengths is depicted in Figure 4, offering a visual representation of the variability in inferred linker lengths across the dataset. Most sequences show lengths consistent with the consensus linker sequence (15 amino acids). Nevertheless, 1,834 scFvs report a linker length of 10 amino acids, warranting special attention and flagging. Other linker lengths display negligible frequencies within the dataset. Nucleotide linker inference is presented in the Supplementary materials (Figure S1, Table S1, Figure S2).

**Table 2.**
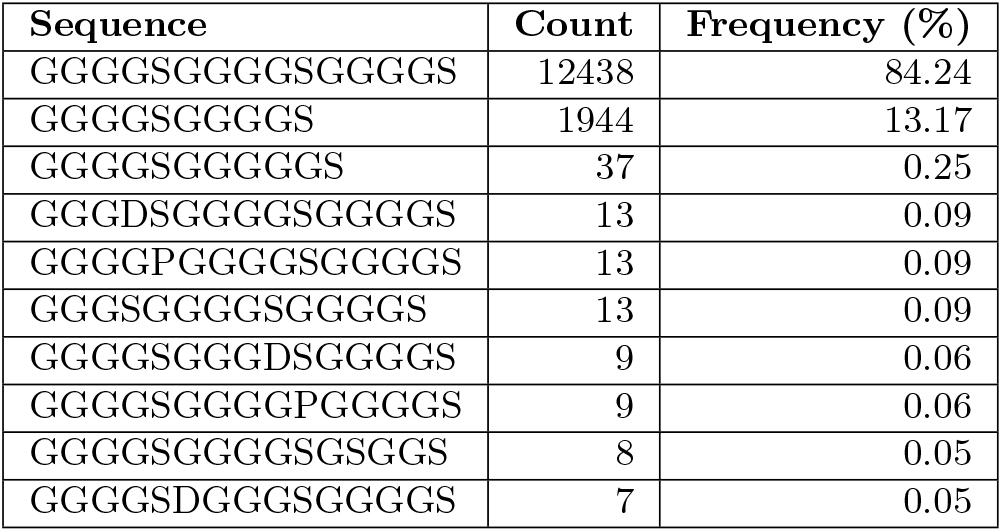
Linker amino acid sequence.

**Figure 3.**
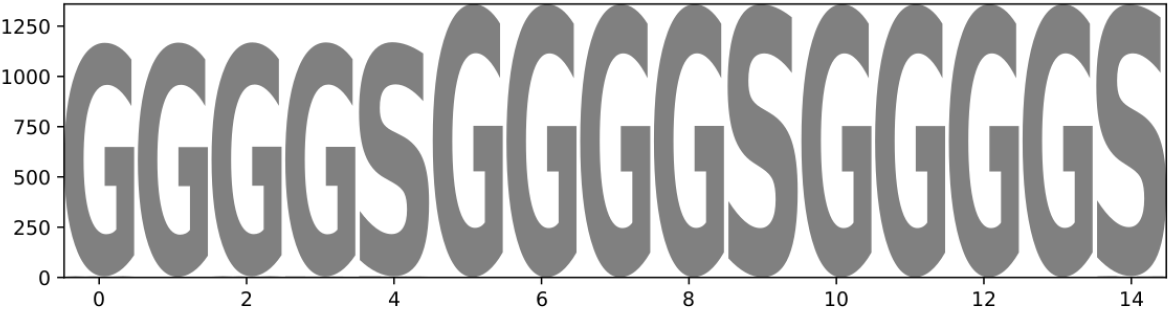
Sequence logo illustrating the amino acid composition at each position of the linker sequence.

**Figure 4.**
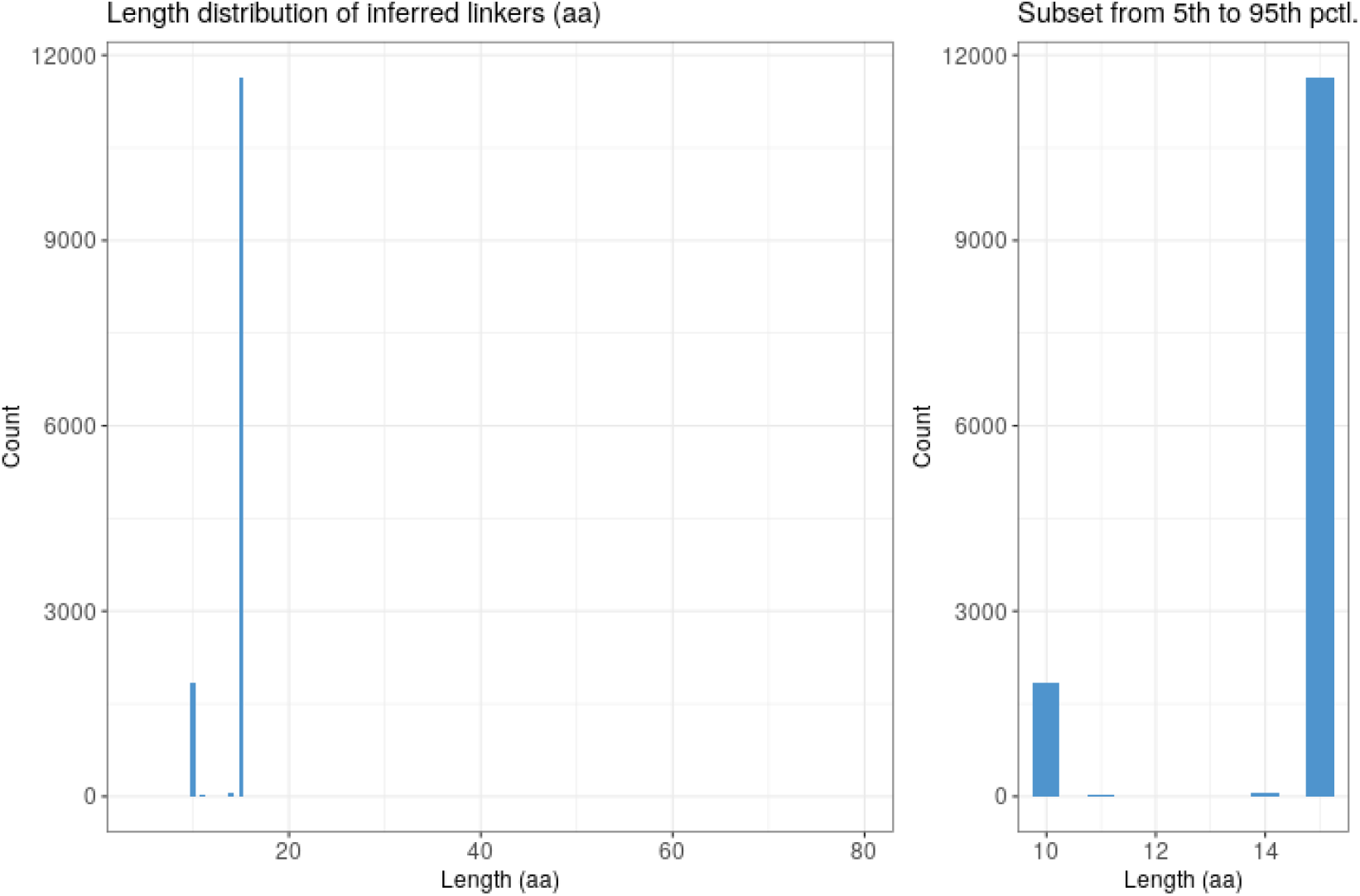
Distribution of inferred amino acid linker lengths within the dataset. Panel A illustrates the distribution across all detected linker lengths, while panel B provides a focused view of the distribution, presenting a histogram within the 5th and 95th percentile interval of linker lengths.

Top 10 most frequent amino acid linker sequences identified within the dataset. Each row corresponds to a unique linker sequence, ranked by its frequency of occurrence. The linker sequence selected as reference is highlighted in bold.

### 3.4 Germline gene usage

One of the advantages of long-read sequencing of scFvs is the ability to evaluate the paired VH-VL germline gene usage comprehensively, which is why we present the chord diagrams generated with R circlize package [16] in Figure 5. These figures provide chord diagram representations of IGHV and IGKV frequency and associations within phage display libraries (no IGLV genes were identified in the dataset). Panel A illustrates the overall number of clones across four panning rounds (0, 1, 2, and 3), where the thickness of the chords reflects the total count of clones associated with each IGHVIGKV combination. In contrast, Panel B focuses on gene family associations, irrespective of clone counts, providing insight into the distinct relationships between IGHV and IGKV gene families. Each chord diagram offers a visual depiction of the distribution of clone counts and gene family associations, shedding light on the dynamics of antibody repertoire diversity within our phage display libraries across multiple panning rounds.

**Figure 5.**
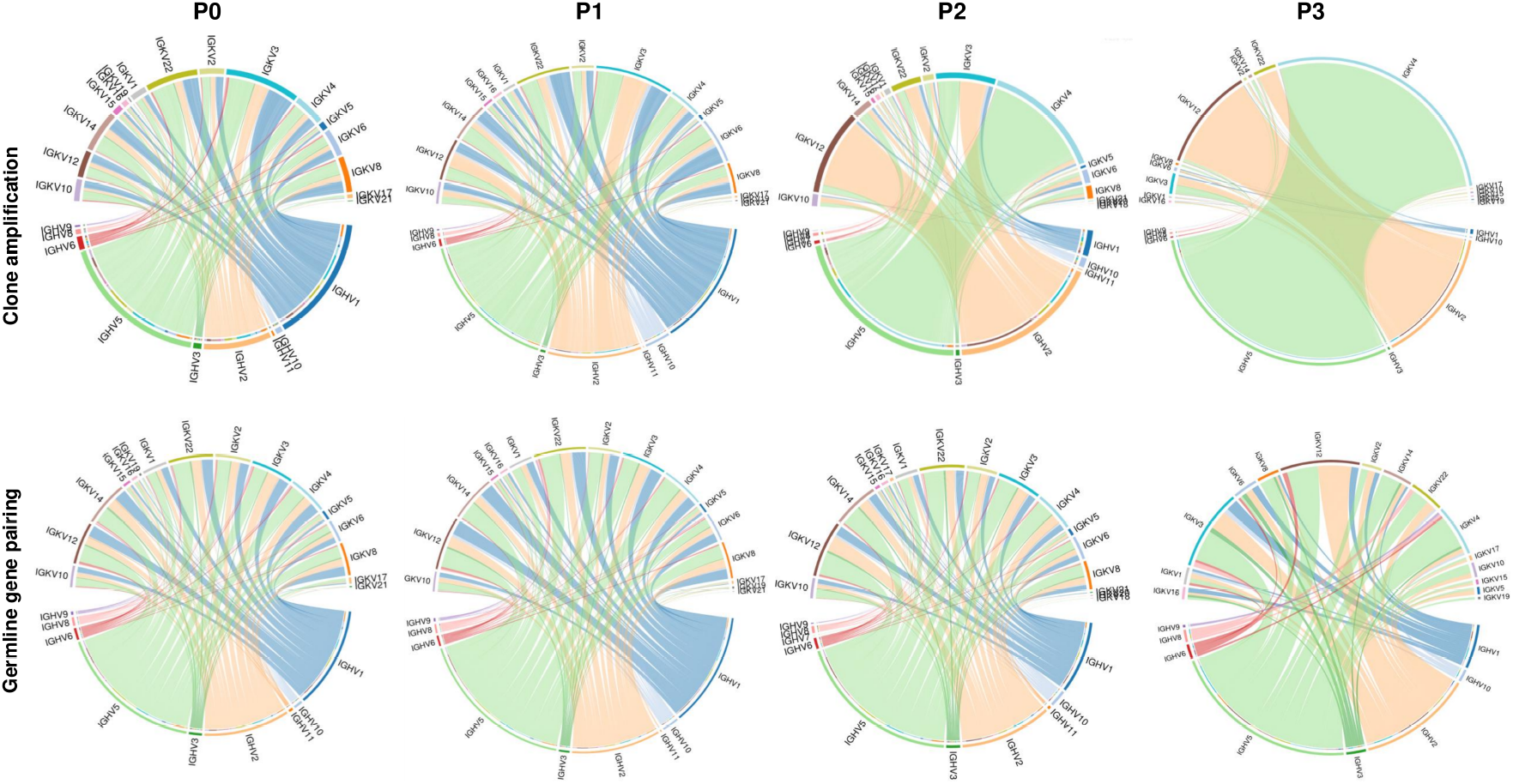
Chord diagram representation of IGHV and IGKV gene family associations.The top pannel depicts overall clone counts across four panning rounds, detailing total occurrences of IGHV-IGKV associations. The bottom pannel illustrates germline gene associations across the same panning rounds, independent of clone counts, showcasing the distinct associations between IGHV and IGKV gene families.

We compared our results with Nannini *et al*. and observed similar trends (Table 3). The percentage of unique heavy and light chain combinations decreased from 68.07% in P0 to 8.27% in P3 in our study, closely mirroring the published dataset which showed a decrease from 67.74% to 16%. The most frequent heavy-light chain combination in P0 was IGHV1-43-IGHJ3 paired with IGKV3S10-IGKJ5, representing 0.55% of our library, while in P3, IGHV5S13-IGHJ2 paired with IGKV4S9-IGKJ2 dominated at 47.43%. Similarly, the published dataset reported the most frequent combination in P0 as IGHV143-IGHJ3 with IGKV3S10-IGKJ5 at 0.36%, and in P3, IGHV5S13-IGHJ2 with IGKV4S9-IGKJ2 at 19.5%. These findings align with the published dataset, leading to the same general conclusion about the selective enrichment of specific scFv clones.

**Table 3.** Heavy and light chain pairing across panning rounds.

| Panning round | Unique VH-VL<br>(% of library) | Most frequent VH-VL |  |  |
| --- | --- | --- | --- | --- |
|  |  | VH | VL | % of library |
| P0 | 68.07 | IGHV1-43-IGHJ3 | IGKV3S10-IGKJ5 | 0.55 |
| P1 | 58.59 | IGHV2-70-IGHJ3 | IGKV3S10-IGKJ5 | 2.73 |
| P2 | 20.81 | IGHV5S13-IGHJ2 | IGKV4S9-IGKJ2 | 20.58 |
| P3 | 8.27 | IGHV5S13-IGHJ2 | IGKV4S9-IGKJ2 | 47.43 |

The differences between Seq2scFv results and those of Nannini *et al*. are expected given the variations in methodology. While Nannini *et al*. used IMGT/HighV-QUEST for their analysis, Seq2scFv employed IgBLAST. Despite utilizing the same germline gene database, the differences in algorithms can lead to variations in gene call assignments. Moreover, the scFv characterization tool in Seq2scFv applies stringent quality control measures: any sequence with poor alignment or containing a stop codons is discarded, regardless of whether these issues occur within the variable domains or the linker sequence. This rigorous approach ensures the high accuracy and reliability Seq2scFv annotations, resulting in a robust characterization of scFvs. Consequently, while the overall trends are consistent with those observed by Nannini *et al*., Seq2scFv’s stringent criteria may account for the minor discrepancies observed between the two datasets.

This table summarizes the percentage of unique heavy and light chain combinations in the scFv library across the panning rounds (P0, P1, P2, P3) and identifies the most frequent heavy-light chain combinations observed at each round. The percentages represent the proportion of the library constituted by these combinations.

## 4 Discussion

NGS guided selection of lead antibody candidates generated by *in vitro* display methods has proven its great potential for accelerating the process of antibody discovery [35] [36] [4]. Moreover, with the progress in high-throughput long-read sequencing, it is now possible to interrogate display libraries at an unparalleled depth coupled with unprecedented levels of detail. Indeed, sequencing of fulllength scFv provides crucial VH-VL pairing information, facilitating a comprehensive evaluation of each candidate’s biophysical properties, germline gene usage, and structure [30] [12]. While it’s very unlikely to sequence 10,000 unique clones through clone picking alone, due to the overrepresentation of dominant clones [12], NGS mining with Seq2scFv of phage display libraries generated by Nannini and colleagues (2020) [30] yielded more than 14,756 high-quality, fully annotated and validated unique scFv sequences. Notably, these libraries were screened using a PacBio RS2 instrument, and new generation PacBio sequencers, such as the Sequel II, offer even greater accuracy and higher throughput, providing larger pools of scFvs to mine.

As the yield of full-length paired antibody data continues to increase, it is crucial for computational tools to keep pace and provide solutions for large-scale comprehensive characterization of scFvs. Seq2scFv, addresses the challenges of characterizing both V-domains in a single read, by combining existing tools for the analysis of single domains (IgBLAST and AntPack) with custom scripts specifically developed to delimit and validate scFvs. Seq2scFv, identified, characterized and validated 17,479 scFvs (representing 14,764 unique scFvs). The results, compiled in a single table, encapsulate the extensive characterization of single-chain variable fragments (scFvs) with a wide array of attributes across several categories. This includes identifiers (such as amino acid and nucleotide SEGUIDs), sequence information (paired VH-VL sequences, scFv and linker start and end positions, translation frames, linker annotations), linker sequence evaluation, quality flags (indicating the validity and productivity of the sequences), and specific annotations for heavy and light chains (covering gene loci, stop codons, frameshifts, and germline alignments). Additionally, the table reports crucial experimental data such as the counts of identical sequences across different panning rounds (P0, P1, P2, P3), enabling the tracking of selection dynamics.

Despite the significant loss of sequences during annotation and scFv validation (74%), it is crucial to emphasize that the retained sequences represent the highest quality subset of scFvs. Indeed, the validated scFv meet significant E-score thresholds for V(D)J alignment and do not present stop codons nor translation frame incongruences between the VH, the VL or the linker sequences. The principal cause of the sequence drop out, accounting for 91% of the discarded sequences, was the presence of stop codons. It is important to also consider that the completeness of the germline gene database will impact the quality and quantity of hits obtained during annotation. This completeness will also aid in identifying the exact coordinates of the V(D)J regions with great accuracy, influencing both the delineation of scFvs and the inference of the linker from the data. Because of this, it is important to continually improve and refine these databases. This is also the reason why the Seq2scFv toolkit allows users to choose their own database or leverage public databases such as OGRDB [21] (available at https://ogrdb.airrcommunity.org/), NCBI (https://ftp.ncbi.nih.gov/blast/executables/igblast/release/database/), or, albeit with licensing restrictions, IMGT ([22]; available at https://www.imgt.org/vquest/refseqh.html). Additionally, Seq2scFv does not remove the scFvs not meeting the quality standards but keeps them in a separate file, allowing users to review and revisit them.

Furthermore, Seq2scFv simplifies downstream analyses such as NGS-guided selection of lead antibody candidates based on frequency. By maintaining a catalog of identical sequences present in each of the sequenced panning rounds, users can track the progressive enrichment of target-binding sequences, which are expected to increase in frequency at later stages of panning. These counts are appended to the final table, enabling users to aggregate the counts at different levels of information. For example, it is possible to identify the enrichment of the scFv at the nucleotide or amino acid levels, or cluster on either the VH or VL, or even any of the CDR or FWR regions. However, in the present article, demonstrating NGS-guided selection of antibody candidates is out of scope.

Seq2scFv’s encoding of sequences also offers a useful solution for future inter-study comparisons. Input reads, nucleotide and amino acid scFvs, are all encoded with a SEGUID, which enables the efficient comparison and identification of identical sequences across panning rounds within the same experiment, or even across different experiments, databases and platforms. Overall, it is a robust feature for tracking and managing antibody data processed with Seq2scFv.

Apart from evaluating annotated scFvs, it is also of great interest to evaluate preprocessing statistics to consider the overall experiment quality. The preprocessing statistics acquired prior to Seq2scFv analyses highlight the number of reads discarded at various filtering stages, including quality, adapter removal, and size filtering, enabling researchers to assess the accurate construction of the libraries. Additionally, by examining read length distributions, researchers can identify sequences at unexpected lengths, uncovering potential issues during library preparation, amplification, or the presence of adapter sequences. Moreover, detecting an increase in the frequency of specific length intervals in later panning rounds implies an effective reduction in antibody diversity and a potential enhancement in specificity as selection progresses, validating the panning process. Sharing feedback with the laboratory based on these observations can enhance experimental practices and improve data quality in subsequent experiments.

In conclusion, Seq2scFv stands as a versatile and accessible toolkit for the analysis of scFvs sequences obtained through long-read sequencing methods. Its open-source nature, free for academic use but subject to a commercial license for commercial purposes, ensures widespread accessibility while addressing proprietary concerns. Notably, the flexibility inherent in Seq2scFv tools enables researchers to customize key parameters such as germline gene databases, numbering schemes, and quality thresholds, thereby tailoring the analytical process to suit specific research objectives. Moreover, the standalone functionality of Seq2scFv provides researchers with the reassurance of data security and privacy, eliminating the need to share sequences online and mitigating potential intellectual property issues. With its emphasis on rapid processing, robust analysis, and comprehensive annotation, we anticipate that Seq2scFv will accelerate the identification and characterization of lead antibody candidates with enhanced efficiency and accuracy.

## 5 Author contributions

MBS developed the toolkit, wrote the manuscript and analyzed the publicly available dataset. LS provided valuable insights on the toolkit’s annotation in relation to its use in downstream analysis. ESS and NL contributed to the processing of the reads used to develop the toolkit. ESS also adapted

*basseto* to the specific requirements of the project. IX provided scientific insights crucial for the design and application of the toolkit. BA played a pivotal role in the development of the tools, provided a general overview of the project, and contributed valuable scientific insights throughout the process. Each author reviewed and approved the final version of the manuscript.

## 6 Disclosure of interest

All authors were employees of JSR Life Sciences at the time the toolkit was developed. The authors declare no other competing financial interests. However, the authors affirm that this potential conflict of interest has not influenced the scientific integrity or objectivity of the research presented in this manuscript.

## 7 Data availability

Seq2scFv code is available on GitHub under https://github.com/ngs-ai-org/seq2scfv. Annotated tables, figures and FASTA files are deposited in Zenodo under 10.5281/zenodo.12635263. Supplementary table 2 is available in the online version of the article.

## 8 Supplementary information

**Figure S1.**
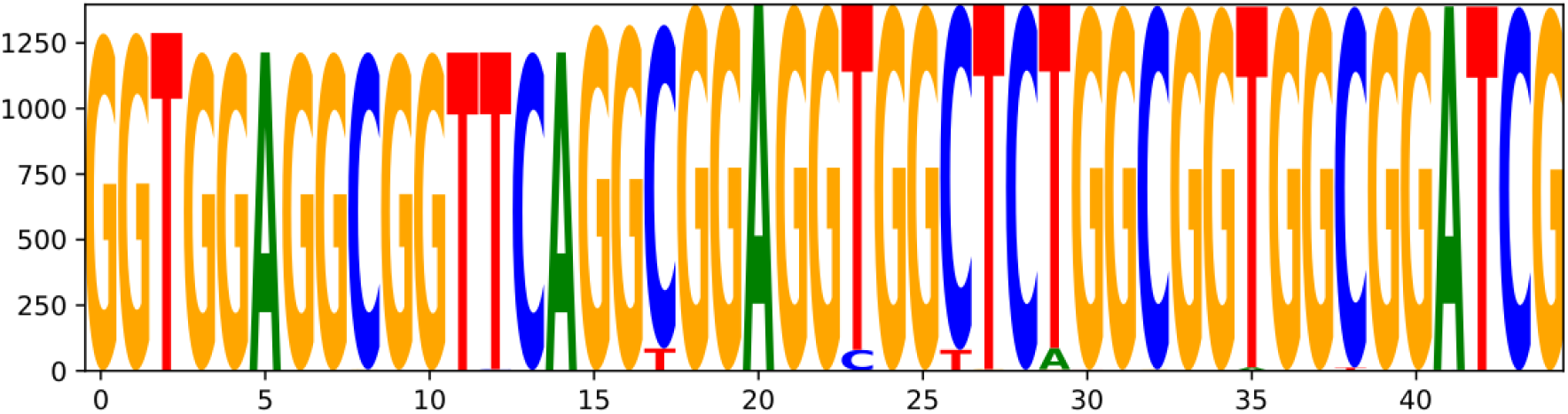
Sequence logo illustrating the nucleotide composition at each position of the linker sequence.

**Table S1.**
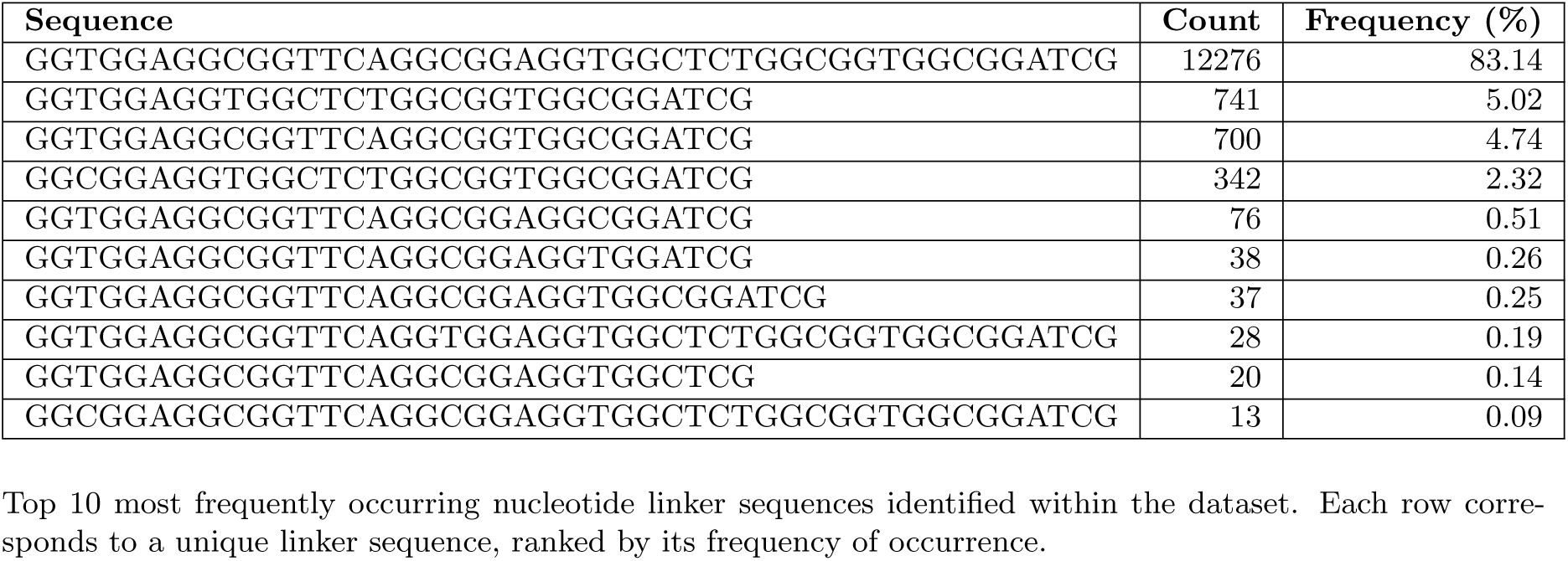
Linker nucleotide sequence.

**Figure S2.**
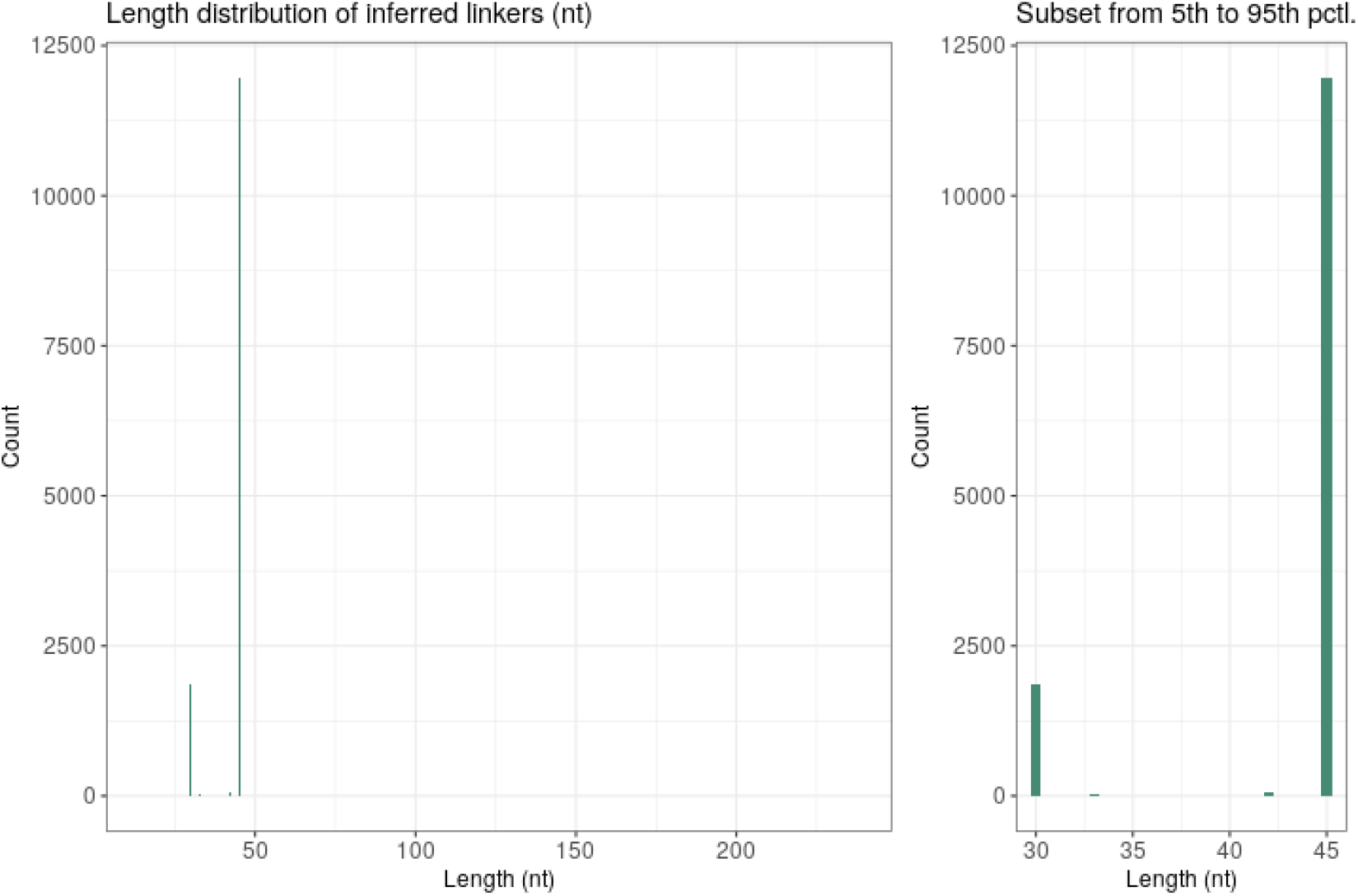
Distribution of inferred nucleotide linker lengths within the dataset. Panel A illustrates the distribution across all detected linker lengths, while panel B provides a focused view of the distribution, presenting a histogram within the 5th and 95th percentile interval of linker lengths.

## Notes

https://github.com/ngs-ai-org/seq2scfv

https://zenodo.org/records/12635263

